# Targeting dermatophyte Cdc42 and Rac GTPase signaling to hinder hyphal elongation and virulence

**DOI:** 10.1101/2024.03.04.583433

**Authors:** Masaki Ishii, Yasuhiko Matsumoto, Tsuyoshi Yamada, Hideko Uga, Toshiaki Katada, Shinya Ohata

## Abstract

The identification of novel molecular targets for antifungal drugs is critical due to limited treatment options and drug-resistance threats. We screened inhibitors of small GTPases, molecular switches in signal transduction, in *Trichophyton rubrum*, the primary cause of dermatophytosis. Our study found that chemical and genetic inhibition of Cdc42 and Rac GTPases, which are involved in cellular morphological changes, significantly impair hyphal formation, and are crucial for pathogenic fungal growth and virulence. Genetic repression of Cdc24, a guanine nucleotide exchange factor of Cdc42 and Rac, led to hyphal growth defects, abnormal cell morphology, and cell death. Chemical screening identified EHop-016 as an inhibitor of Cdc24 activity, which improved outcomes in *in vitro* nail infection and invertible infection models of *T. rubrum*. Our results suggest the Cdc24-Cdc42/Rac pathway as a promising therapeutic target for antifungal agent development, with EHop-016 as a potential lead compound.

## INTRODUCTION

Dermatophytes, a type of common infectious fungus, are responsible for dermatophytosis, a condition that affects over 100 million people worldwide.^1–4^ Despite the existence of antifungal treatments, dermatophytosis often proves to be stubborn and recurrent, particularly in the case of tinea unguium, a type of nail infection,^5^caused by dermatophytes. The dermatophytes have the ability to invade superficial host tissues such as hair and skin and, in rare instances, can infect dermal tissues and lymph nodes in immunocompromised patients.^6^ Conditions such as dermatophytosis and onychomycosis can lead to severe asthma and foot ulcers in patients with diabetes mellitus.^7, 8^ The current antifungal drugs target a limited number of fungal enzymes involved in the synthesis of ergosterol, such as squalene epoxidase (allylamine and benzylamine classes) and sterol 14α-demethylase (azole class).^9^ However, the emergence of drug-resistant strains underscores the urgent need for the discovery of new therapeutic targets for dermatophytes.^5,10,11^

In order to establish an infection, fungal spores adhere to the superficial layer of the host’s skin, germinate, form hyphae through cell-proliferation and morphological elongation, and penetrate the superficial tissue. Pathological observations, reconstructed skin experiments, and *ex vivo* analysis suggest that germination and hyphal growth are critical steps in colonization and infection.^12, 13^ In *Trichophyton rubrum*, the most common cause of dermatophytosis,^1^ previous studies have identified genes related to drug-resistance and pH- response.^10,14–17^ However, due to a lack of genetic techniques, the genes involved in hyphal growth in *T. rubrum* remain largely unidentified.

Small GTPases function as molecular switches in intracellular signaling, altering their conformation based on the guanine nucleotide forms they bind.^18^ The guanosine diphosphate (GDP)-binding (inactive) form is exchanged for the guanosine triphosphate (GTP)-binding (active) form by the Dbl-homology (DH) and pleckstrin-homology (PH) domains (hereafter collectively referred to as the DH-PH domain) of guanine nucleotide exchange factors (GEFs). Small GTPases play a key role in germination and hyphal growth in human pathogenic fungi.^19,20^ For instance, Rac1 and Rac2 are synthetically essential for hyphal growth in *Cryptococcus neoformans*, and Arl1 is a crucial protein for invasive filamentous growth in *Candida albicans*. The Rho family, a well-studied and conserved small GTPase family, activates its effectors to regulate cell morphology and growth by modulating actin organization.^21^ In *Saccharomyces cerevisiae*, the Rho family small GTPase Cdc42 is vital for growth and viability,^22–24^ whereas Cdc42 and another Rho family protein Rac are synthetic lethal in the filamentous fungi *Ustilago maydis*, *Aspergillus nidulans*, and *Aspergillus niger*.^25–27^ In *Neurospora crassa* and *Claviceps purpurea*, the DH-PH domain-containing protein Cdc24 activates Cdc42 and Rac.^28, 29^ Since Cdc24 is an essential protein and a modulator of hyphal formation in some fungi,^28–32^ the activation machinery of Cdc42 and Rac by Cdc24 could be a novel therapeutic target for the development of antifungal agents. However, the function of these proteins in dermatophytes is unknown, and chemical compounds that inhibit GEF-mediated fungal Rac and Cdc42 activation have not been identified.

In this study, we established a method to suppress gene expression in *T. rubrum* in a copper-dependent manner. We discovered that both chemical and genetic inhibition of the Cdc24-Rac/Cdc42 signaling pathway significantly impedes germination and hyphal formation. EHop-016, an inhibitor of the interaction between Cdc24 and Cdc42/Rac, improved *in vitro* nail infection and reduced the lethality of silkworms injected with *T. rubrum*. Our results propose that targeting small GTPase signaling, especially the Cdc24- Rac/Cdc42 pathway, holds promise as a therapeutic approach for developing antifungal agents. Additionally, EHop-106 could serve as a lead compound.

## RESULTS

### Mammalian Rac and Rac/Cdc42 inhibitors suppress the germination of *T. rubrum*

To investigate a small GTPase pathway involved in germination and mycelial growth in *T. rubrum*, we screened small GTPase inhibitors *in cellulo* by culturing fungal conidia with them. Seven compounds known to inhibit mammalian Rho and Arf family proteins—Rac (EHT1864 and NSC23766), Rac and Cdc42 (AZA1), Rho A (Rhosin and Y16), and Arf (Brefeldin A and NAV2729) were utilized.^33–38^ EHT1864, NSC23766, and AZA1 (Figure 1A) hindered germination and mycelial growth in *T. rubrum*, while Rhosin, Y16, Brefeldin A, and NAV2729 did not (Figure 1B, C). These Rac and Cdc42 inhibitors suppressed germination in a dose-dependent manner (Figure 1D). The ED_50_ of EHT1864, NSC23766, and AZA1 were 16 µM, 640 µM, and 127 µM, respectively (Table 1). These findings imply that Rac and/or Cdc42 signaling is crucial for germination and hyphal growth in *T. rubrum,* suggesting these pathways as potential therapeutic targets for dermatophytosis.

**Figure 1:**
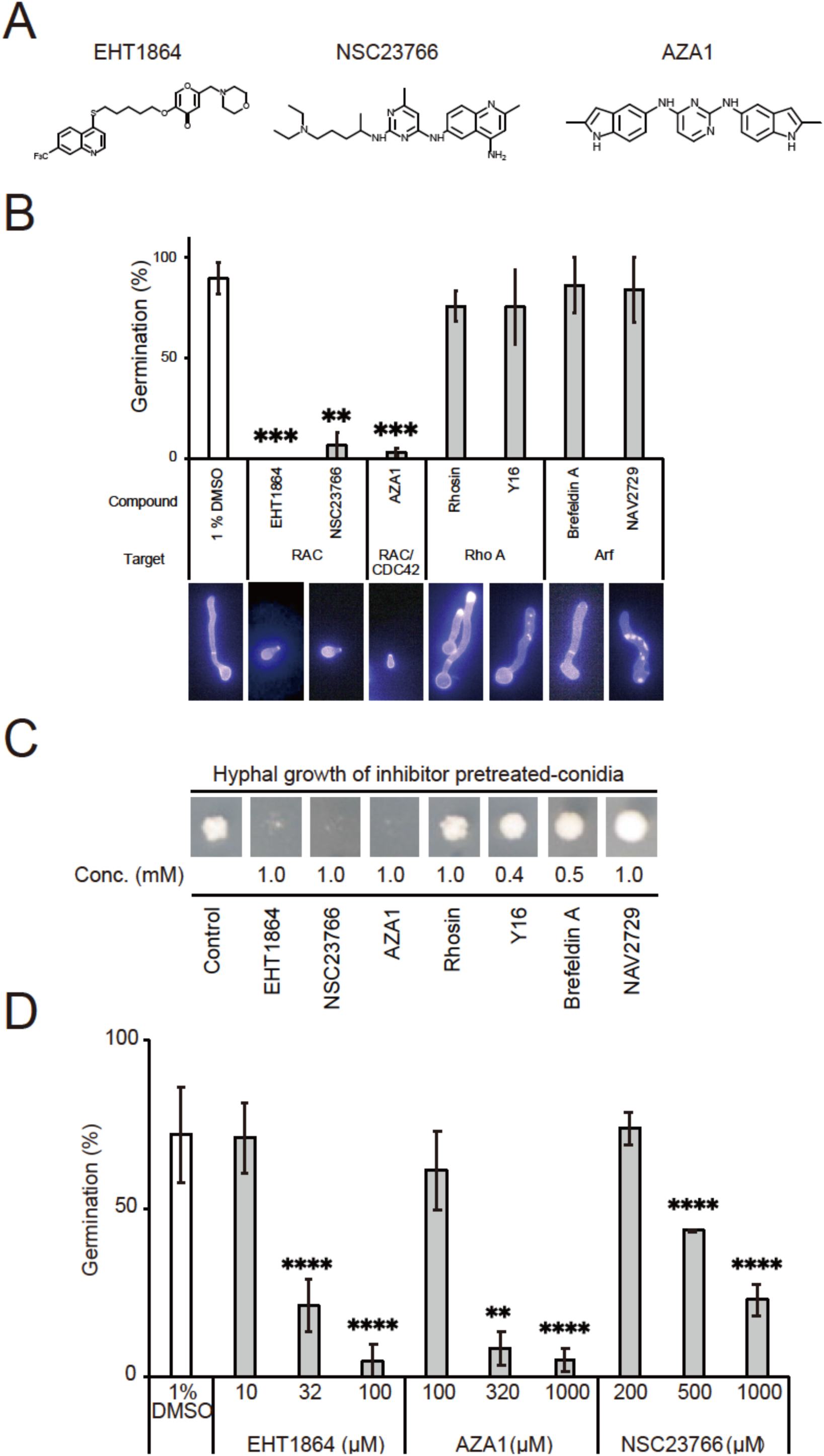
Mammalian Rac and Rac/Cdc42 inhibitors suppress *Trichophyton rubrum* germination. A. Structures of Rac inhibitors EHT1864, NSC23766, and Rac/Cdc42 inhibitor AZA1. B. Conidia were incubated with 1% DMSO (control) or mammalian small GTPase inhibitors at 28°C overnight, stained with calcofluor white, and observed under fluorescent microscopy for quantification of conidia and germinated conidia. n = 3 each. **P < 0.01; ***P < 0.001. Mean ± SD. C. Conidia were incubated with 1% DMSO (control) or mammalian small GTPase inhibitors at 28°C overnight, washed with saline, and incubated on agar for 4 days to observe fungal growth. D. Conidia were incubated with indicated concentrations of mammalian Rac and Rac/Cdc42 inhibitors at 28°C overnight, stained with calcofluor white, and observed under fluorescent microscopy for quantification of conidia and germinated conidia. n = 3 each. **P < 0.01; ****P < 0.0001. Mean ± SD.

**Table 1.** Effect of mammalian Cdc42 and/or Rac inhibitors against fungal Cdc42, Rac, and germination in *T. rubrum*.

| Inhibitor | Target in mammalian cell | TrCdc42 IC<sub>50</sub><br>(μM) | TrRac IC<sub>50</sub><br>(μM) | IC<sub>50</sub> to<br>*T. rubrum*<br>germination<br>(μM) |
| --- | --- | --- | --- | --- |
| AZA1 | Rac1 and Cdc42 | 147 | 127 | 127 |
| CASIN | Intersectin1/Cdc42 | 192 | 518 | 20 |
| Ehop-016 | Vav/Rac | 40 | 137 | 45 |
| EHT1864 | Rac1 | 154 | >1000 | 16 |
| ITX3 | TrioN/Rac, Rho | >50 | >50 | >50 |
| Ketorolac | Rac1, Cdc42 | 889 | 1416 | 363 |
| MBQ-167 | Rac, Cdc42 | >100 | >100 | >100 |
| ML-141 | Cdc42 | >1000 | >1000 | 1.7 |
| NSC23766 | Tiam-1, Trio/Rac | 368 | 477 | 640 |
| ZCL278 | Intersectin1/Cdc42 | >1000 | >1000 | >1000 |

### XP_003237709 and XP_003234353 are *T. rubrum* homologs of Rac and Cdc42, respectively

Rac and Rac/Cdc42 inhibitors fall into two categories: one inhibiting the binding of Rac/Cdc42 to guanine nucleotides, e.g., EHT1864,^39^and the other inhibiting the binding of Rac/Cdc42 to their GEF, e.g., NSC23766. ^35^ As both types of inhibitors inhibited hyphal growth of *T. rubrum* (Figure 1), we focused on identifying Rac and Cdc42 homologs in *T. rubrum*. Through a BLAST search in the genome sequence of the *T. rubrum* CBS118892 strain,^40^we identified XP_003237709 and XP_003234353 of *T. rubrum* CBS118892 as candidate homologs of human Rac1 (identity: 80%) and Cdc42 (identity: 78%). Phylogenetic tree analysis demonstrated that XP_003237709 and XP_003234353 are related to Rac and Cdc42 proteins, respectively, from other species (Figure 2A). We expressed recombinant XP_003237709 and hexahistidine (His)-tagged XP_003234353 proteins (Figures 2C and E). An ammonium sulfate accelerates guanine nucleotide association and dissociation of GTPases.^41^ These proteins bound to Bodipy-labeled GTP in an ammonium sulfate-dependent manner (Figure 2B, D, and F). These findings affirm that XP_003237709 and XP_003234353 are *T. rubrum* homologs of Rac and Cdc42, subsequently referred to as TrRac and TrCdc42, respectively.

**Figure 2:**
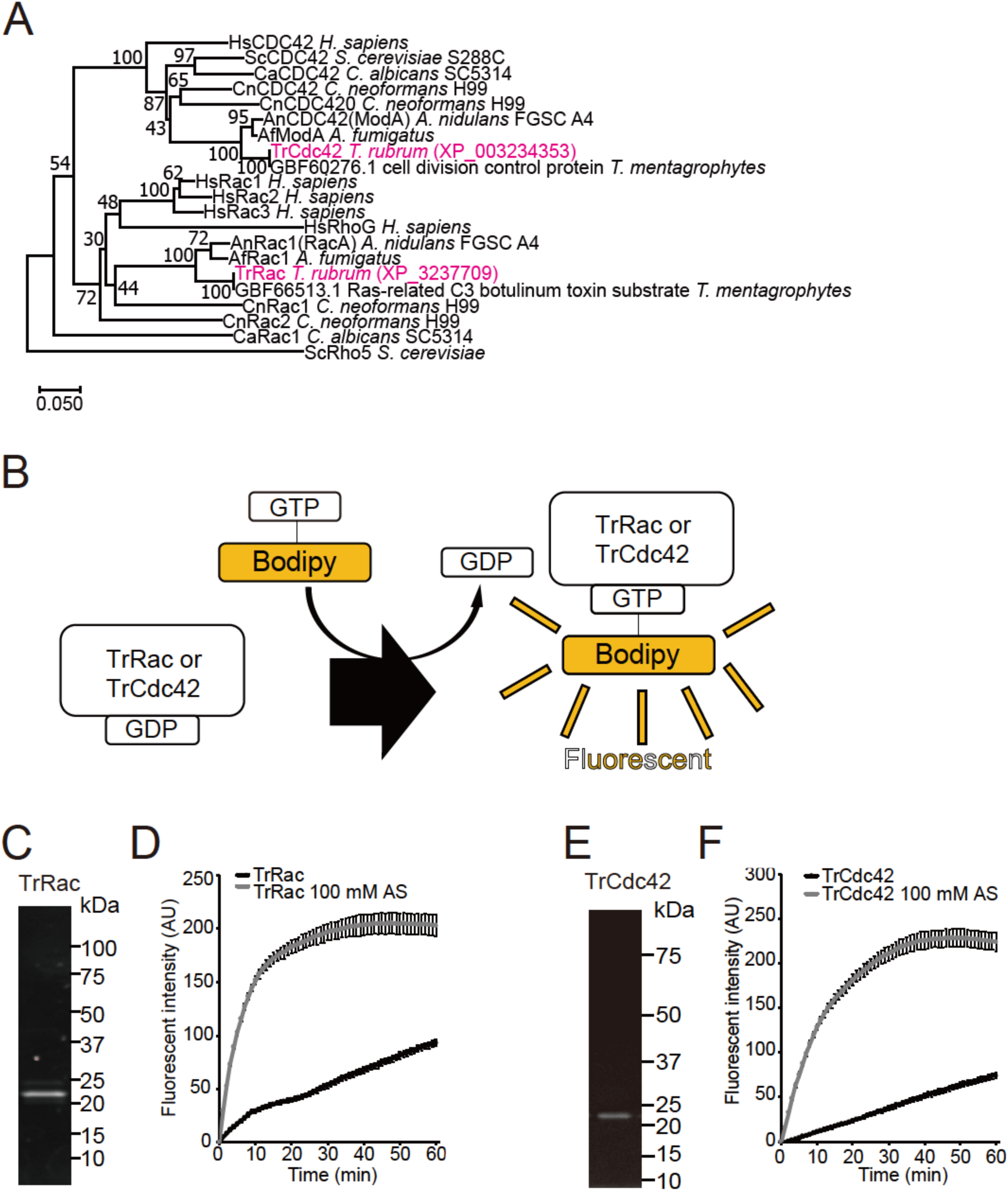
XP_003237709.1 and XP_003234353.1 are TrRac and TrCdc42 in *Trichophyton rubrum*. A. Phylogenetic tree of fungal Rac and Cdc42 proteins using the neighbor-joining method. The percentage of replicate trees clustering taxa together in bootstrap test (500 replicates) is indicated at branches. Evolutionary distances were calculated using Poisson correction method in terms of amino acid substitutions per site. B. Fluorescence-based guanine nucleotide exchange assay method. C. SDS-PAGE analysis of purified recombinant TrRac proteins stained with Stain-Free fluorescence stain. D. TrRac guanine nucleotide exchange assay was performed. One μM recombinant TrRac and 0.1 μM Bodipy GTP were mixed with or without 100 mM ammonium sulfate (AS), and fluorescence intensity was measured. Mean ± SD. E. SDS-PAGE analysis of purified recombinant TrCdc42-His proteins stained with Stain-Free fluorescence stain. F. TrCdc42-His guanine nucleotide exchange assay was performed. One μM recombinant TrCdc42-His and 0.1 μM Bodipy GTP were mixed with or without 100 mM ammonium sulfate (AS), and fluorescence intensity was measured. Mean ± SD.

### *T. rubrum* Cdc24 promotes guanine nucleotide exchange of TrRac and TrCdc42

In the filamentous fungi *N. crassa* and *C. purpurea*, DH-PH domain-containing proteins act as GEFs, i.e., these proteins promote the conversion of Rho-type GTPases, such as Rac and Cdc42, from GDP-bound to GTP-bound forms.^28,29^ We searched the *T. rubrum* genome for DH-PH domain-containing proteins and found XP_003232406. We cloned this cDNA, but the size of the PCR product (registered in GenBank as MW699015) was 402 bp shorter than the predicted size from XP_003232406. Sequence alignment analysis revealed that the last part of the third exon of XP_003232406.1 is missing in MW699015 (Figure S1A). As this sequence is also missing in Cdc24 cDNAs from other filamentous fungi (Figure S1B), we assumed that this additional sequence in XP_003232406 is due to misprediction of the splice site. The sequence similarity of the protein encoded by MW699015 to *S. cerevisiae* Cdc24 was 27%. Phylogenetic tree analysis revealed that MW699015 and Cdc24 proteins from other fungi belong to the same clade (Figure 3A). We therefore designated the protein encoded by MW699015 as TrCdc24. A purified recombinant peptide containing DH-PH domains (from 150^th^ to 508^th^ amino acids in Fig. 3B) expressed in E. coli (Figure 3B and C) enhanced the guanine nucleotide exchanges of TrRac and TrCdc42- His (Figure 3D and E). These results suggest that TrCdc24 is a GEF for TrRac and TrCdc42.

**Figure 3:**
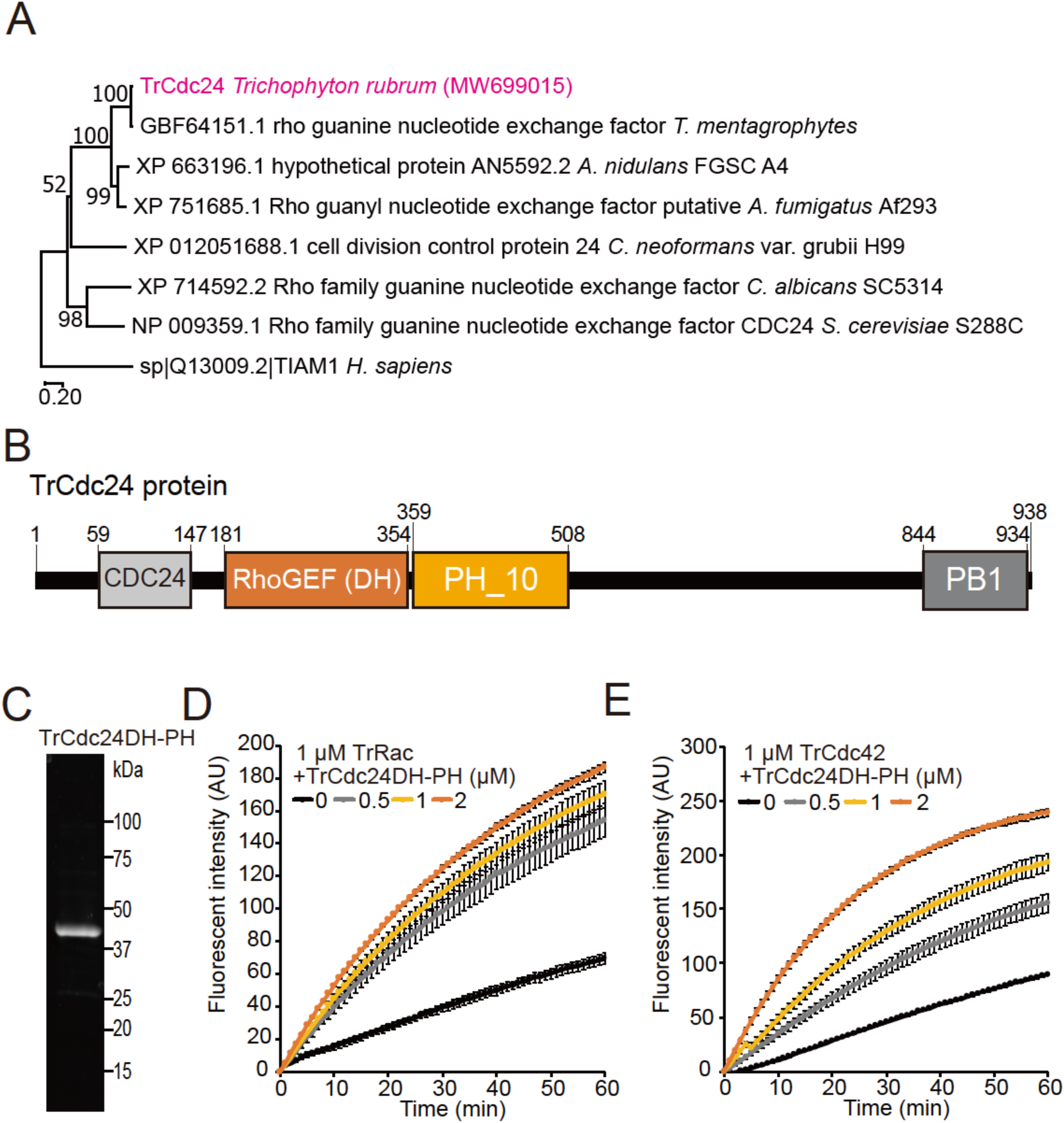
*Trichophyton rubrum* Cdc24 stimulates guanine nucleotide exchange of Rac and Cdc42. A. Phylogenetic tree analysis of fungal Cdc24 and human Tiam1 using the neighbor-joining method. The percentage of replicate trees clustering taxa together in bootstrap test (500 replicates) is indicated at branches. Evolutionary distances were calculated using Poisson correction method in terms of amino acid substitutions per site. B. Protein domains of Cdc24 shown in an image. C. SDS-PAGE analysis of purified recombinant TrCdc24 proteins stained with Stain-Free fluorescent stain. D. Guanine nucleotide exchange of TrRac with or without TrCdc24 was observed. One μM TrRac and 0.1 μM Bodipy GTP were incubated with or without 0.5, 1, and 2 μM TrCdc24DH-PH. Mean ± SD E. Guanine nucleotide exchange of TrCdc42-His with or without TrCdc24 was observed. One μM TrCdc42-His and 0.1 μM Bodipy GTP were incubated with or without 0.5, 1, and 2 μM TrCdc24DH-PH. Mean ± SD.

### Repression of Tr*cdc24* leads to inhibition of mycelial growth

As homokaryotic knockout of Cdc24 is lethal in *N. crassa*,^28^ it is anticipated that TrCdc24 is similarly indispensable for survival in *T. rubrum*, preventing us from elucidating the function of TrCdc24 in hyphal growth in a TrCdc24 knockout strain. To circumvent the potential lethality associated with a TrCdc24 knockout, we sought to establish a conditional gene repression system. We utilized a copper-responsive promoter (P*_ctr4_*), a system employed in another dermatophyte, *Trichophyton mentagrophytes (formerly Arthroderma vanbreuseghemii),* to repress gene expression in a copper concentration-dependent manner.^42^ We integrated P*_ctr4_*upstream of Tr*cdc24*’s open reading frame (ORF) (Figure 4A) and confirmed its integration through Southern blot analysis (Figure 4B). The expression level of Tr*cdc24* mRNA was significantly reduced in P*_ctr4_*Tr*cdc24* mutant *T. rubrum* cultured with copper ions, depending on the copper ion concentration, compared to that in wild type (WT) and P*_ctr4_*Tr*cdc24* mutant *T. rubrum* cultured with the copper chelator bathocuproinedisulfonic acid (BCS) (Figure 4C). This copper ion concentration-dependent decrease in Tr*cdc24* mRNA expression was not observed in WT (Figure 4C). These results suggest that the P*_ctr4_*system functions effectively in *T. rubrum*. Intriguingly, the P*_ctr4_*Trc*dc24* mutant experienced arrested mycelial growth under gene silencing conditions in a copper ion concentration-dependent manner (Figure 4D), highlighting the essential role of TrCdc24 in *T. rubrum* mycelial growth.

**Figure 4:**
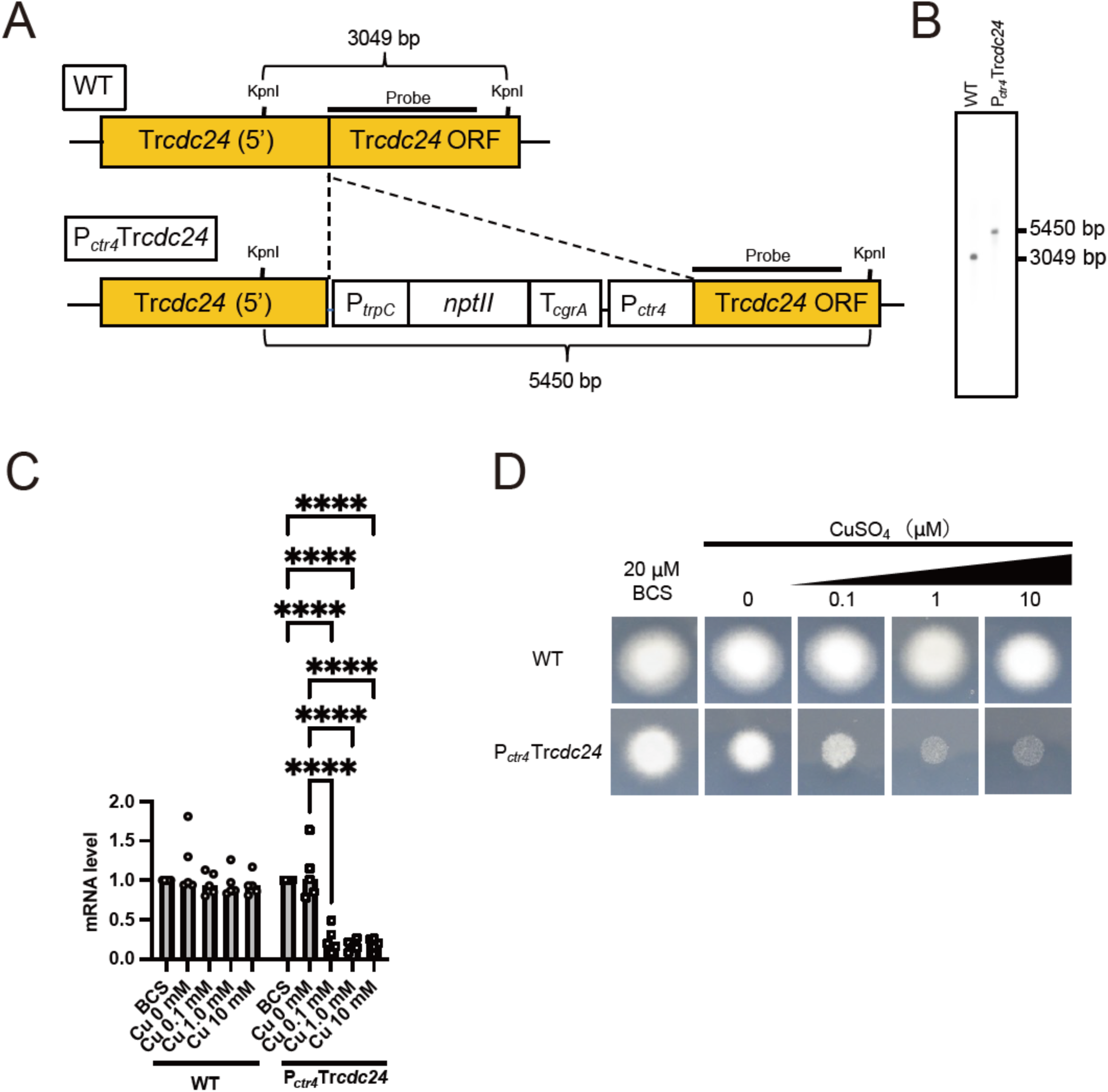
Repression of the Tr*cdc24* gene shows mycelial growth inhibition. A. Schematic representation of the Tr*cdc24* locus in the genome of *Trichophyton rubrum* CBS118892 wild type and P*_ctr4_*Tr*cdc24* mutant. B. Southern blot analysis of genome DNA samples from *T. rubrum* CBS118892 wild type and P*_ctr4_*Tr*cdc24*. C. Quantification of Tr*cdc24* mRNA in total RNA of *T. rubrum* CBS118892 and P*_ctr4_*Tr*cdc24* cultured in medium containing 0, 0.1, 1, or 10 μM CuSO_4_ or 20 μM Bathocuproinedisulfonic acid (BCS), a copper ion chelator. n = 5 each. ****P < 0.0001. Mean ± SD. D. Mycelial growth of *T. rubrum* CBS 118892 and P *_ctr4_*Tr*cdc24* on an agar plate with 0, 0.1, 1, or 10 μM CuSO_4_or 20 μM BCS.

### TrCdc42 is essential for mycelial growth in a TrRac deletion background

Since mammalian Rac inhibitors inhibit germination and hyphal formation in *T. rubrum* (Figure 1B-D) and repression of TrRac GEF TrCdc24 caused a defect in mycelial growth (Figure 4D), we generated a conditional Tr*rac* suppression mutant (P*_ctr4_*Tr*rac*) and observed mycelial growth. Contrary to our expectation, no difference in mycelial growth was observed between the fungi under induced and suppressed gene expression conditions in the P*_ctr4_*Tr*rac* mutant (data not shown). Furthermore, we were able to knock out the Tr*rac*. The Tr*rac* gene deletion and selection-marker insertion in this strain were confirmed by PCR (Figure 5A-C). Furthermore, the mRNA level of Tr*rac* in the ΔTr*rac* strain was confirmed to be reduced to a level comparable to the limit of detection (data not shown). No apparent defect in mycelial growth was observed in the strain obtained (Figure 5D). Since the amino acid sequence of TrCdc42 is similar to that of TrRac and the guanine nucleotides bound to TrCdc42 were exchanged by TrCdc24 as to TrRac, we hypothesized that TrCdc42 acts in concert with TrRac in mycelial growth. Therefore, we investigated the inhibitory effect of Rac inhibitors on the guanine nucleotide exchange reaction of TrCdc42 by TrCdc24. TrCdc42 guanine nucleotide exchange by TrCdc24 was inhibited by the Rac inhibitors EHT1864 and NSC23766 at 154 and 368 μM, respectively (Table 1). To further investigate the function of TrCdc42 in mycelial growth, we generated a conditional Tr*cdc42* repression mutant (P*_ctr4_*Tr*cdc42*) and a ΔTr*rac* and P*_ctr4_*Tr*cdc42* double mutant and confirmed the insertion of the fragment, which is containing selective marker and *ctr4* promoter, into targeted region by PCR (Figure 5E, F). In P*_ctr4_*Tr*cdc42* strain, Tr*cdc42* mRNA level was decreased under CuSO_4_ treatment (Figure 5G). P*_ctr4_*Tr*cdc42* showed a slight growth defect under repressive conditions, whereas this growth defect was enhanced in the Tr*rac*- deficient/P*_ctr4_*Tr*cdc42* strain under repressive conditions (Figure 5D). These results suggest that TrRac and TrCdc42 have similar essential roles in mycelial growth and that the TrCdc24-TrCdc42/TrRac pathway is essential for mycelial growth.

**Figure 5:**
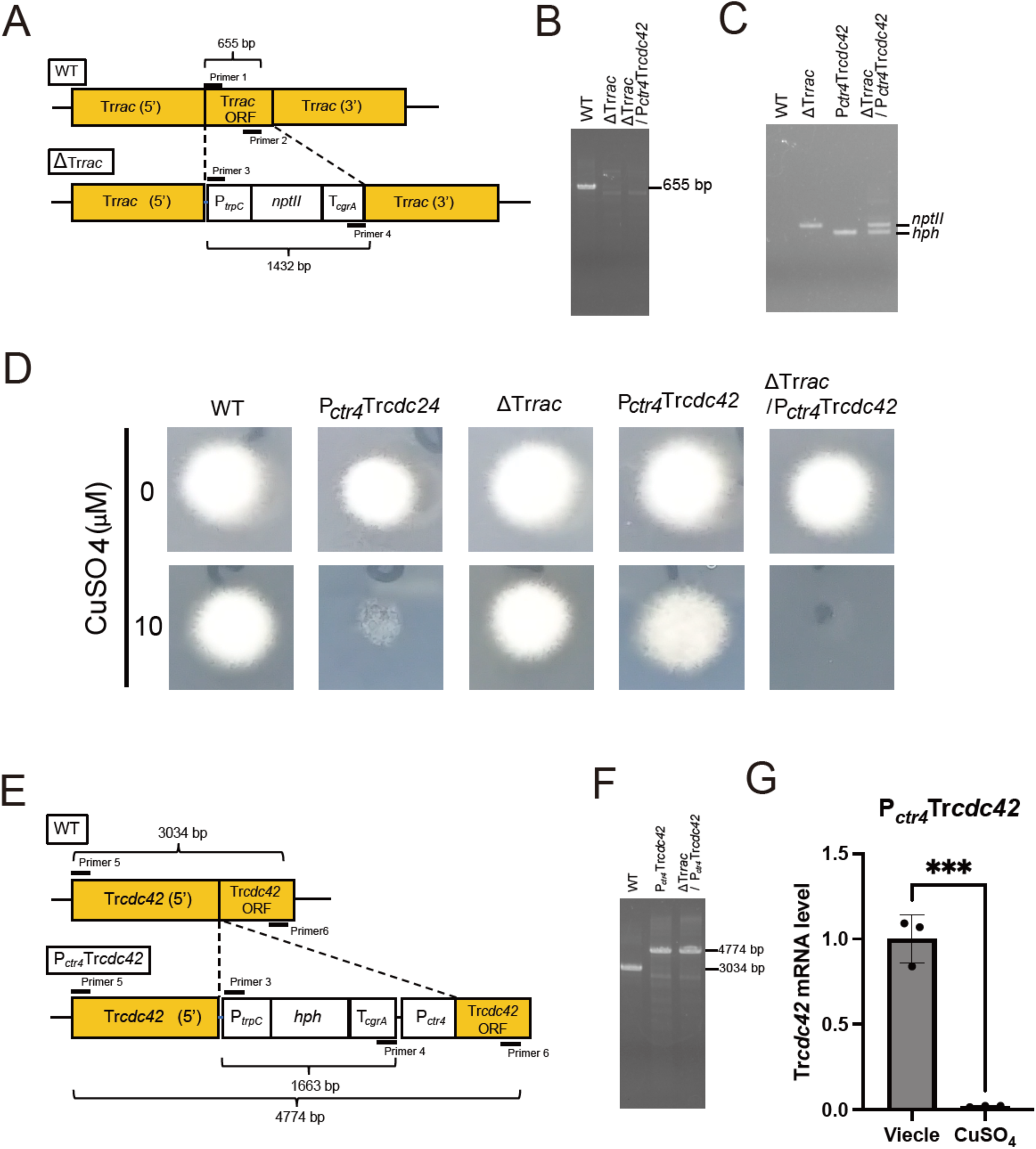
Repression of the Tr*cdc42* gene shows mycelial growth inhibition in a TrRac deletion background. A. Schematic representation of the Tr*rac* locus in the genome of *Trichophyton rubrum* CBS118892 WT and Tr*rac* deletion mutant (ΔTr*rac*). B. PCR analysis conducted on genomic DNA samples from *T. rubrum* CBS118892 WT, P*_ctr4_*Tr*raci*, and ΔTr*rac*/ P *_ctr4_*Tr*cdc42* utilizing primers 1 and 2 as depicted in panel A. C. PCR analysis was conducted on genomic DNA samples from *T. rubrum* CBS118892 WT, P*_ctr4_*Tr*raci*, P *_ctr4_*Tr*cdc42*, and ΔTr*rac*/ P *_ctr4_*Tr*cdc42* utilizing primers 3 and 4 as depicted in panel A. D. Mycelial growth of *T. rubrum* CBS 118892, P *_ctr4_*Tr*cdc24*, ΔTr*rac*, P *_ctr4_*Tr*cdc42*, ΔTr*rac*/ P *_ctr4_*Tr*cdc42* on an agar plate with 0 or 10 μM CuSO_4_. E. Schematic representation of the Tr*cdc42* locus in the genome of *T. rubrum* CBS118892 wild type and P*_ctr4_*Tr*cdc42* mutant. F. PCR analysis was conducted on genomic DNA samples from *T. rubrum* CBS118892 WT, P *_ctr4_*Tr*cdc42*, and ΔTr*rac*/ P *_ctr4_*Tr*cdc42* utilizing primers 5 and 6 as depicted in panel A. H. Quantification of Tr*cdc42* mRNA in total RNA of *T. rubrum* CBS 118892 and P*_ctr4_*Tr*cdc42* cultured in medium containing 0 or 10 μM CuSO_4_. n = 3 each. ***P < 0.001. Mean ± SD.

### Repression of TrCdc24 leads to abnormal cell morphology, actin delocalization and cell death

To gain insights into the halted mycelial growth in *T. rubrum* when the TrCdc24- TrRac/TrCdc42 pathway is repressed, we conducted a chronological observation of the morphology of P*_ctr4_*Tr*cdc24* (Figure 6A). In the absence of copper ions, oval cells observed on day 1 elongated into filamentous cells on day 2. This morphological elongation between culture days 1 and 2 was evident when calculating the polarity index (major axis length/minor axis length) (Figure 6B). On the contrary, the P*_ctr4_*Tr*cdc24* mutant cultured with copper ions for two or three days exhibited a more rounded morphology, characterized by a shorter major axis and a longer minor axis compared to the mutant cells cultured without copper ions (Figure 6A). The polarity index of the copper-treated cells was significantly lower than that of the untreated cells (Figure 6B). Interestingly, by day 3, the cell size of the copper-treated cells showed no significant difference from that of the untreated cells (Figure 6C). These findings indicate that hyphal formation is regulated by TrCdc24.

**Figure 6:**
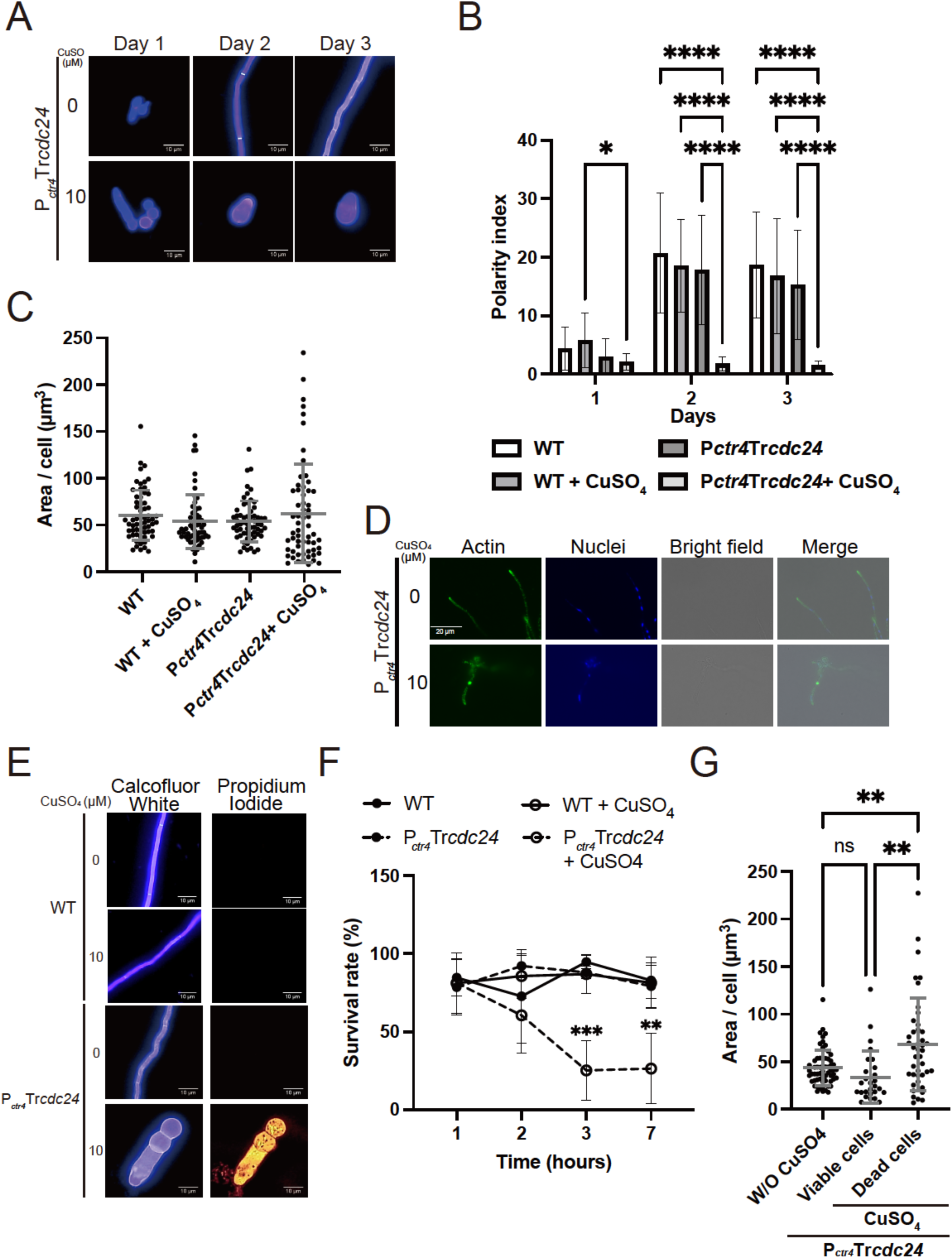
Repression of the Tr*cdc24* gene shows abnormal cell formation, actin localization, cell death, and reduced cell viability. A. P *_ctr4_*Tr*cdc24* conidia (2 x 10^6^) were cultured in RPMI medium with or without 10 μM CuSO_4_ for 3 days. Fungal cells were collected on days 1, 2, and 3, stained with calcofluor white, and observed under a fluorescence microscope. B. WT and P *_ctr4_*Tr*cdc24* conidia (2 x 10^6^) were cultured in RPMI medium with or without 10 μM CuSO_4_ for 3 days. Fungal cells were collected on days 1, 2, and 3, stained with calcofluor white, and observed under a fluorescence microscope. The polarity index was calculated on days 1(n = 50–60 each), and 2 (n = 50–60 each), and day 3 (n = 60 each). * P < 0.05. ****P < 0.0001. Mean ± S.D. C. Cell area was measured on day 3. The dots on the graph represent the value of the individual cell area. n = 60 each. Mean ± SD. D. Actin was detected with an anti-actin antibody and Alexa 488-conjugated anti-mouse IgG (green). Nuclear DNA was stained with DAPI (blue). E. P *_ctr4_*Tr*cdc24* conidia (2 x 10^6^) were cultured in RPMI medium with or without 10 μM CuSO_4_ for 7 days. Fungal cells were stained with calcofluor white and PI. The stained cells were observed under a fluorescence microscope. F. WT and P *_ctr4_*Tr*cdc24* conidia (2 × 10^6^) were cultured in RPMI medium with or without 10 μM CuSO_4_ for 7 days. The fungal cells were sampled on days 1, 2, 3, and 7 and stained with calcofluor white and observed by fluorescence microscopy. n = 3 each. **P < 0.01. ***P < 0.001. Mean ± SD. G. P *_ctr4_*Tr*cdc24* conidia (2 x 10^6^) were cultured in RPMI medium with (CuSO_4_) or without (W/O CuSO_4_) 10 μM CuSO_4_ for 7 days. The area of cells was measured on day 7. The dots on the graph represent the value of the individual cell area. The numbers of W/O CuSO_4_, viable and dead cells were 60, 27, and 42, respectively. **P < 0.01. Mean ± SD.

Both Rac and Cdc42 play roles in facilitating actin organization.^21^ In filamentous fungi, F-actin localizes to the hyphal tip and forms higher-order structures known as actin patches and actin cables.^43^These structures are responsible for endocytosis and polarized transport, respectively. In the absence of copper ions, actin was localized at the hyphal tip (Figure 6D). Conversely, Tr*cdc24* repression impaired actin localization at the hyphal tip, resulting in abnormal actin accumulation in the cytosol (Figure 6D). This leads to the conclusion that TrCdc24 regulates the localization of actin at the hyphal tip.

In yeast, actin dysfunction leads to cell death.^44^ To elucidate whether TrCdc24 functions in cell death, dead cells were stained with propidium iodide (PI), a dyepermeating dead cells but not living cells. Under the repression condition, P*_ctr4_*Tr*cdc24* mutants were stained with PI after culturing for three days (Figure 6E). The survival rate of Tr*cdc24*-repressed cells was reduced to 22% after three days of culture (Figure 6F). To further analyze the relationship between abnormalities in actin-regulated cell morphology and cell death, we compared the size of living and dead cells under Tr*cdc24*-suppressed conditions. The results showed an increase in cell size in dead cells (Figure 6G). These findings suggest that TrCdc24 contributes to mycelial growth by modulating cell morphology and viability.

### A TrCdc24 inhibitor EHop-016 ameliorates *in vitro* nail infection and lethality in an invertebrate infection model of dermatophytosis

Given the implications from the above results, suggesting that the activation mechanism of TrRac and TrCdc42 is a promising target for antifungal drug development, we screened using Rac and Cdc42 inhibitors previously reported in mammals, including Ehop-016, AZA1, EHT1864, CASIN, NSC23766, ketorolac, MBQ-167, ML-141, ZCL278, and ITX3.^45^ Employing an *in vitro* biochemical assay, we evaluated the impact of Rac/Cdc42 inhibitors on the potentiating effect of TrCdc24 in the GDP/GTP exchange reaction of TrRac and TrCdc42. The IC_50_ of these inhibitors against TrRac and TrCdc42-His exceeded 127 μM and 40 μM, respectively (Table 1). Notably, in this biochemical screen, EHop-016 exhibited the strongest activity against TrCdc42, with an IC_50_ of 40 µM (Figure 7A, B, and Table 1). Those inhibitors showing TrCdc42 inhibitory activity in the biochemical assay also inhibited germination *in celluro* (Figure 7C and Table 1). Intriguingly, the IC_50_ of these inhibitors against TrCdc42 showed a higher correlation coefficient with fungal germination inhibition compared to that against TrRac (0.58 and 0.39, respectively). These findings suggest that TrCdc42 is likely to play a more crucial role than TrRac in *T. rubrum* germination, consistent with the observations that the P*_ctr4_*Tr*cdc42* strain exhibited partially reduced mycelial growth under gene repression conditions, while the P*_ctr4_*TrRac and Rac deletion strains displayed no change in mycelial growth under all tested conditions (Figure 5A).

**Figure 7:**
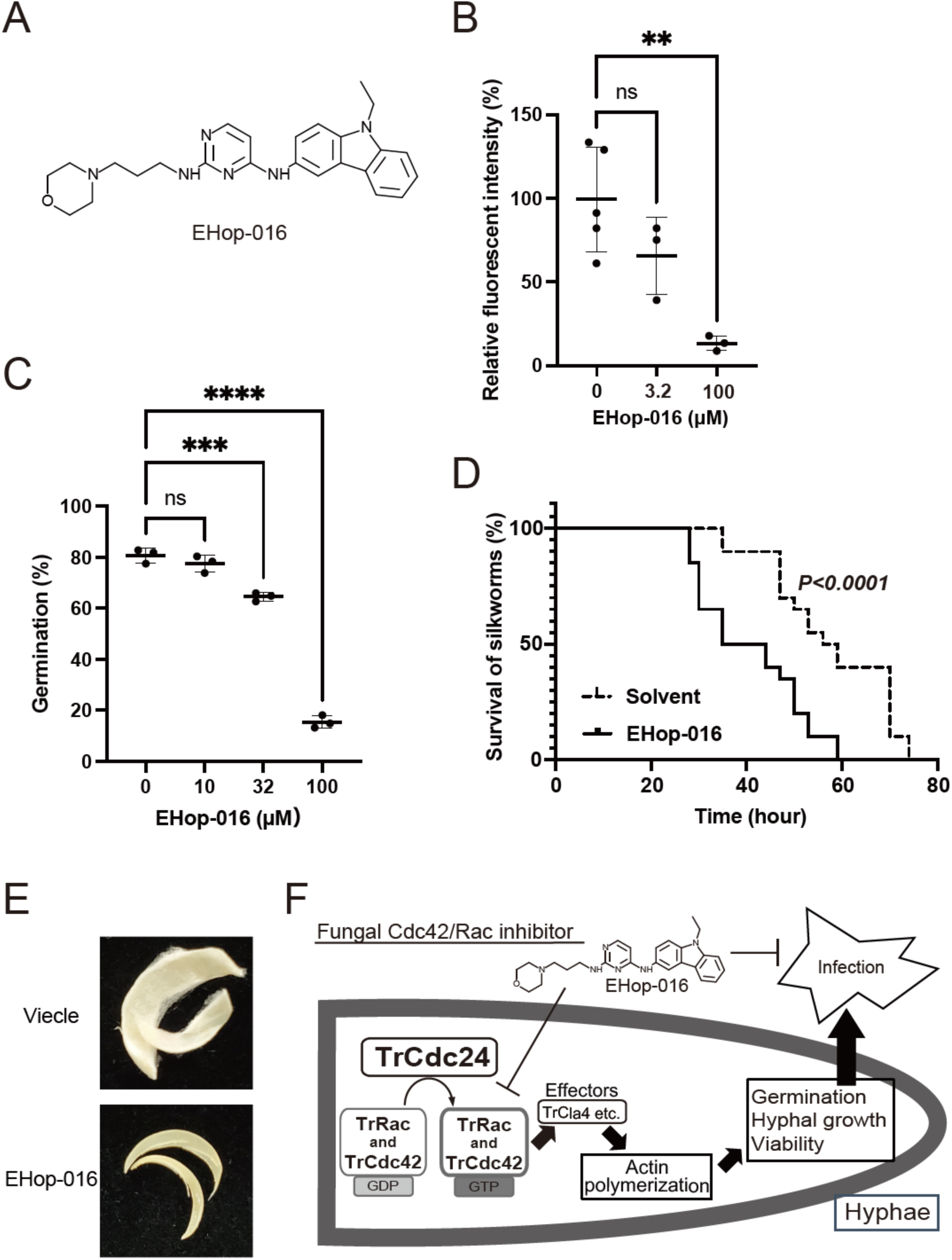
Therapeutic efficacy of EHop-016 against an invertebrate infection model of dermatophytosis. A. Structure of the fungal Cdc42 and Rac inhibitor Ehop-016. B. One μM TrCdc42-His and 0.1 μM Bodipy GTP were incubated with 1 μM TrCdc24DH- PH and 0 (n = 5), 32 (n = 3), 100 (n = 3) μM EHop-016. The fluorescence intensity was measured after 1 hour. **P < 0.01. Mean ± S.D. C. Conidia were incubated with the indicated concentration of EHop-016 at 28°C overnight and then stained with calcofluor white. Conidia and germinated conidia were observed under fluorescence microscopy for quantification. n = 3 each. ***P < 0.001. ****P < 0.0001. Mean ± SD. D. Therapeutic effect of EHop-016 in a silkworm infection assay with *T. rubrum*. Conidia (1 x 10^8^/ml) of *Trichophyton rubrum* were injected into the silkworm, followed by 10 mM of EHop-016 or vehicle were injected into the silkworm. The survival of silkworms was observed. E. Conidia (1 x 10^4^) of *T. rubrum* were spread on a nail, followed by 60 mM of EHop-016 or vehicle, and applied to the nail every day for four days. After 36 days of incubation at 28°C, the nails were observed. F. Model. TrCdc42 and TrRac are activated to their GTP-bound forms by TrCdc24. Subsequently, the activated GTPases bind to effector molecules, like TrCla4, to enhance the polymerization of actin. This, in turn, promotes germination, hyphal growth, and viability, ultimately leading to an enhanced infection by the fungus. The employment of a fungal Cdc42 and Rac inhibitor, EHop-016, demonstrated the potential to mitigate the fungal infection.

Given that *T. rubrum* is an anthropophilic but not zoophilic dermatophyte, establishing an *in vivo* infection model in vertebrates is challenging.^46^To determine the efficacy of chemical inhibition of the Rac/Cdc42 pathway in animal infection, we utilized silkworms, a well-established invertebrate infection model for various fungi, including dermatophytes.^48–50^ EHop-016 was selected as the test compound due to its higher TrCdc42 inhibitory activity, and its pharmacological parameters had already been reported in animal.^47^ EHop-016 significantly extended the survival of animals infected with dermatophytes (Figure 7D), indicating that an inhibitor targeting the Rac/ Cdc42 pathway assists animals in recovering from dermatophyte infection.

Onychomycosis, primarily caused by *T. rubrum*, represents a challenging and recurring nail infection.^48^ To elucidate the effect of the Rac/Cdc42 inhibitor against *T. rubrum* nail infection, we tested EHop-016 in an *in vitro T. rubrum* nail infection model. Treatment with 2.6% EHop-016 resulted in lesser *T. rubrum* growth on the nails compared to the vehicle treatment control (Figure 7E), confirming that the Rac/Cdc42 inhibitor EHop-016 inhibits *T. rubrum* hyphal growth on nails.

## DISCUSSION

In the realm of antifungal chemotherapy, significant challenges arise from the limited pool of target molecules and the emergence of chemoresistant fungi against existing drugs.^49^ Through the development of a gene silencing system in *T. rubrum*, we made two pivotal discoveries. First, we identified the TrCdc24-TrRac/TrCdc42 pathway as a promising therapeutic target for dermatophytosis. Second, we pinpointed chemical inhibitors that target this pathway. Our study demonstrated that TrCdc24 serves as a GEF for TrRac/TrCdc42 and is indispensable for mycelial growth, cell morphogenesis, and viability. Inhibitors of Rac/Cdc42 curtail the activation of TrRac and TrCdc42 by TrCdc24, subsequently impeding the germination of *T. rubrum*. EHop-016, the most potent Cdc42 inhibitor in this study, showcased the potential to ameliorate the lethality observed in silkworms injected with *T. rubrum*, a well-established *in vivo* model for dermatophyte infection.^50^ This study underscores the possibility that inhibiting the TrCdc42-TrRac/TrCdc42 signaling pathway could serve as an effective therapeutic strategy against fungal infections.

### Function of TrCdc24-TrRac/TrCdc42 pathway

In both animals and fungi, Rac and Cdc42 play pivotal roles in actin organization.^21^ This study demonstrates that TrCdc24 enhances the guanine nucleotide exchange reaction of TrRac and TrCdc42 (Figure 3) and reveals that repression of TrCdc24 expression impacts hyphal formation, actin localization at the hyphal tip, and cell viability. Biochemical, genetic, and chemical-biological analyses suggest that TrRac and TrCdc42 proteins also contribute to mycelial growth in *T. rubrum*. The regulation of actin localization by TrCdc24 is likely mediated by the activation of TrRac and TrCdc42. Recently, we have reported that TrCla4 is a PAK in *T. rubrum* that is activated by TrRac and may regulate mycelial growth by enhancing actin polymerization.^51^ The TrCDC24-TrCDC42/TrRac-TrCla4 pathway may promote hyphal growth by regulating the actin dynamics (Figure 7F). While the downstream signaling of TrRac and TrCdc42 is not fully clarified, studies from other filamentous fungi and yeast suggest that the downstream signal may be mediated not only by PAKs but also by effector molecules that regulate actin polymerization, such as Formin.^26,52^ In a yeast actin mutant strain with reduced ATPase activity, cell death was observed along with abnormal localization of actin filament and subsequent loss of actin filament.^53^ Similar cell death was observed in the TrCdc24-suppressed strain with abnormal actin localization, suggesting that control of actin localization may also contribute to survival in dermatophytes.

### Potential of antifungal drug discovery by targeting Cdc42 and Rac: its generality and specificity

Cdc42 and Rac play a significant role in cell migration and defense against external stimuli in human skin.^54–56^ Furthermore, inhibition of Pak2, an effector of CDC42 and Rac, may lead to acute cardiovascular toxicity.^57^ Thus, the use of TrCdc42 and TrRac inhibitors for the treatment of dermatophytosis could potentially cause adverse effects such as immunosuppression, infections, and acute cardiovascular toxicity through the inhibition of human CDC42, Rac, and/or off-targets. These side effects could be avoided due to the low amino acid sequence identity (∼27%) between human Rho GEFs and *T. rubrum* Cdc24. Therefore, the Cdc24-mediated Rac/Cdc42 activation machinery is a promising target for antifungal drug development. The germination inhibition assay revealed that MBQ-167, a potent mammalian Rac and Cdc42 GEF interaction inhibitor, did not show any inhibitory activity against the fungus even at 100 μM.^58^ This suggests that the compounds can recognize differences between mammalian and fungal p21 (Rac and Cdc42). Calcineurin, another signaling molecule that controls hyphal growth, has been proposed as a target molecule for antifungal agent development.^59^ Although FK506 (tacrolimus), an approved immunosuppressant and inhibitor of calcineurin, inhibits fungal growth *in vitro*,^60,61^, its immunosuppressive effect is a deterrent for an antifungal drug. To overcome this problem, derivatization has been attempted, and some derivatives have been successful in reducing the immunosuppressive action despite having strong antifungal activity.^61,62^ In addition, the crystal structure of calcineurin-FK506-FKBP12 was recently elucidated and was used for the design of novel fungal-specific FK506 analogs.^63^ Structural analysis and derivatization of lead compounds such as EHop-016 are expected to be powerful strategies for the discovery of fungal selective Rac/Cdc42 inhibitors.

The amino acid sequences of the Cdc24 GEF exhibit striking similarity among filamentous fungi, with a notable 73% identity to the DH-PH domain of *A. fumigatus* Cdc24. Rac and/or Cdc42 play pivotal roles in hyphal growth in pathogenic fungi like *A. niger* and *C. albicans,* where Cdc24 is identified as an indispensable protein for hyphal growth in *C. albicans*.^26,32^ Considering these findings, it is highly plausible that inhibitors targeting the TrCdc24-TrRac/TrCdc42 pathway could serve as broad-spectrum antifungal drugs.

### The difference between animal and fungal targets of Rac and Cdc42 inhibitors

Rac and Cdc42 have garnered attention as potential anticancer drug targets, leading to the discovery of over 30 mammalian Rac and/or Cdc42 inhibitors.^45^ Our chemical inhibitor screening revealed a notable difference in selectivity between mammalian and fungal Rac/Cdc42. Specifically, the mammalian Rac-selective inhibitors, EHT1864, and EHop-016 showed higher selectivity for Cdc42 than for Rac in fungi (Table1). This finding challenges the conventional notion that mammalian Rac inhibitors selectively inhibit fungal Rac and Cdc42, akin to their selectivity in mammals. The mechanism underlying this selectivity of inhibitors remains unclear, and this discovery provides valuable insights for understanding the function of fungal Rac and Cdc42 using these inhibitors.

### A genetic tool to analyze genes of interest

*T. rubrum* stands as the most common causative fungus of dermatophytosis;^1^ however, the molecular mechanism governing hyphal growth remains unclear. A prior histological study suggested that hyphal growth is essential for *T. rubrum* virulence,^12^ but limited genetic tools and the slow growth of this fungus have prevented further analysis. Several studies of genetic modification in this fungus have been reported.^14,15,17,64^ Recently, auxotrophic mutants of *T. rubrum* have been generated using the Cas9 system.^65^ In these articles, gene deletions and gene insertions were performed for functional analysis. Previous methods have made it difficult to analyze the functions of essential proteins that are potential drug targets. In the present study, we implemented a copper-responsive promoter system previously utilized in *Arthroderma vanbreuseghemii,*^42, 60^ enabling us to explore essential gene functions and exercise spatiotemporal regulation of gene expression. This system is poised to streamline the investigation of *T. rubrum* pathogenicity and the identification of novel drug targets.

## Conclusion

We have demonstrated that the chemical and genetic inhibition of the TrCdc24- TrRac/TrCdc42 pathway leads to growth arrest in *T. rubrum*. In this study, we proposed an approach that combines genetic and biochemical methods to discover novel antifungal compounds against *T. rubrum*. It is anticipated that screening for more potent and selective inhibitors of the fungal Cdc24-Cdc42/Rac pathway will yield novel antifungal agents. This study presents Cdc42-Rac/Cdc42 signaling as a novel drug target, distinct from those of existing antifungal drugs.

### Significance (284/300 words, no reference)

Superficial fungal infections, such as athlete’s foot, impact over 10% of the world’s population, significantly affecting their quality of life. Deep fungal infections like aspergillosis can often be fatal. The number of patients affected by these infections continues to rise due to an aging population and advances in immunosuppressive treatments. Treatment-resistant fungi pose a concern, and there are limited antifungal drug targets. In *T. rubrum*, the most common anthropophilic dermatophyte worldwide, we focused on the small GTPase pathways that act as molecular switches in signal transduction. Given the limited genetic techniques for *T. rubrum*, we successfully adapted a method to repress gene expression in the presence of copper ions. This technology enables the analysis of previously inaccessible essential genes, contributing to the search for new drug targets and the functional analysis of these genes. We found that the fungal Rac/Cdc42 GTPase pathway promotes mycelial growth in this fungus. Fungal Cdc24, an activator of Rac/CDC42 GTPases, has little similarity to human homologous proteins, making it a promising antifungal target. We identified EHop-016 as a fungal Rac/Cdc42 inhibitor by screening known mammalian Rac/Cdc42 inhibitor. EHop-016 not only inhibits dermatophyte germination but also showed therapeutic activity in an invertebrate model of dermatophyte infection and inhibited fungal growth on the human nail. EHop-016 can be used to analyze the molecular functions of other pathogenic fungi, such as *Aspergillus fumigatus* and *Rhizopus oryzae*, which cause life-threatening infections. In conclusion, by developing a new genetic technique to evaluate candidate antifungal drug targets in *T. rubrum*, this study has identified fungal Rac/Cdc42 as a potential antifungal target and EHop-016 as a promising lead antifungal compound.

## Acknowledgment

We thank Emiri Horikoshi, Sayumi Takahashi, Rika Matsubara, Konatsu Yoshida, and Tsuyoshi Ojima for their technical assistace. This work was supported by JSPS KAKENHI Grant Numbers 16H06163, 17H03981, 19K16656, 20K06550, 20K07038, 21K15438, and 23K06533 and Takeda Science Foundation.

## Author Contributions

Conceptualization, M.I., T.K., and S.O.; Methodology, T.Y.; Investigation, M.I., H.U., and Y.M.; Writing – Original Draft, M.I., T.K., and S.O.; Writing – Review & Editing, M.I., T.Y., H.U., Y.M., T.K. and S.O.; Funding Acquisition, M.I., T.K., and S.O.; Resources, T.Y.; Supervision, T.K., and S.O.

## Declaration of Generative AI and AI-assisted technologies in the writing process

No use of AI and AI-assisted technologies.

## STAR Methods

### Bacterial and fungal strains and culture conditions

*T. rubrum* CBS118892, *Escherichia coli* BL21, and *Agrobacterium tumefaciens* EAT105 were utilized. ^66–68^ *E. coli* BL21 was cultured in LB medium with appropriate antibiotics at 37°C. *A. tumefaciens* was cultured in LB or *Agrobacterium* induction medium supplemented with 0.2 mM acetosyringone at 28°C.^69^ *T. rubrum* was cultured at 28°C. Conidia of *T. rubrum* were prepared using a previously described method.^70^ Sabouraud agar or RPMI with MOPS agar, was used for hyphal formation. To evaluate mycelial growth inhibition by inhibitors, a modified 1/10 Sabouraud glucose agar (containing Bacto peptone 0.2%, glucose 0.1%, KH_2_PO_4_ 0.1%, MgSO_4_ · 7H_2_O 0.1%, Bacto agar 1.5%, pH unadjusted) was employed.

### Inhibitors

NSC23766 and MBQ-167 were procured from Selleck Chemicals, USA. EHT1864, ML- 141, EHop-016, ZCL-278, CASIN, Ketorolac, and Brefeldin A were obtained from Cayman Chemical Company, Germany. AZA1, Rhosin, ITX3, and Y16 were sourced from Merck, Germany. NAV2729 was acquired from TOCRIS Bioscience, UK. All inhibitors were solubilized in DMSO.

### Germination assay

Fungal conidia (2 × 10^6^) were subjected to an overnight incubation in Sabouraud medium, supplemented with 1% DMSO (control) or the desired concentration of Rac and/ or Cdc42 inhibitors. Following the incubation, the fungal conidia were washed twice with saline and then stained with 0.2 mM calcofluor white. The germinated and ungerminated conidia were subsequently observed and counted under a fluorescent microscope (Olympus BX53). For the purpose of this experiment conidia, were considered germinated if they exhibited a protrusion that exceeded the diameter of the conidia.

## Plasmid construction

Tr*rac*, Tr*cdc42*, and Tr*cdc24*DH-PH were cloned into pGEX6p-1 using the In-Fusion system (TaKaRa Bio). The amplified DNAs were inserted into the EcoRI-BamHI sites of the pGEX6p-1 using synthesized primer pairs (5′- GGGGCCCCTGGGATCCATGGCTTCTGGTCCAGCTAC -3′ and 5′-<colcnt=3> GTCGACCCGGGAATTCTACTTTCTCTTTGGTTTCTGAGTTGGA −-3′, 5′- GGGGCCCCTGGGATCCATGGTTGTTGCTACTATCAAGTGTG-3′ and 5′- GGAATTCCGGGGATCCTACAAGAGTAGACAGCGGCTC and 5′- GGGGCCCCTGGGATCACTCGAGCTTCAAGGCCAAC -3′ and 5′- GTCGACCCGGGAATTCCGCTGCTTGTCGATATCCT-3′, respectively) were synthesized. To prepare TrCdc42-His, TrCdc42 containing pGEX6p-1 was used as a template and a specific primer pair (F: 5′- CATCACCATCACCATCACTGACGATCTGCCTCGC-3′ and R: 5′- ATGGTGATGGTGATGCAAGAGTAGACAGCGGCTCG−3′) was used. These sequences were sequenced post-cloning.

A Tr*cdc24*-targeting vector, pAg1P*_ctr4_*Tr*cdc24*, was constructed as follows: Approximately 1.8 and 1.9 kb of the upstream and ORF fragments of Tr*cdc24* (TERG_02186) were amplified from *T. rubrum* total DNA by PCR with specific primer pairs. (F: 5′-GGGAAACGACAATCTATAACCCCGGTGATGGAGGA-3′ and R: 5′-TCAATATCATCTTCTACCCCCAATAGCCAAACTGG-3′; F2: 5′-TACAAAGCCTGCGAAATGGAGGGAATGAACGGGAA−3′ and R2: 5′-GTGAATTCGAGCTCGTGGAAACTTGCTGGTCTGGT−3′, respectively).

The basic structure of pAg1^71^ and Pctr4 cassette (P*trpC* [GenBank accession no. X02390], nptII, T*cgrA* [AFUA_8G02750], and P*ctr4* [TERG_01401]) were amplified from pAg 1−3′- UTR of ARB_02021 by PCR with specific primer pairs (F: 5′- CGAGCTCGAATTCACTGGCC−3′ and R: 5′-AGATTGTCGTTTCCCGCCTT−3′; F2: 5′-AGAAGATGATATTGAAGGAGCA−3′ and R2: 5′-TTCGCAGGCTTTGTACTTT−3′, respectively).

These four amplified fragments were joined with the In-Fusion system (TaKaRa Bio).

A Tr*rac*-targeting vector, pAg1ΔTr*rac*, was constructed similarly: Approximately 1.6 and 1.0Lkb of the upstream and downstream fragments of Tr*rac* (TERG_02424) were amplified from *T. rubrum* total DNA by PCR with specific primer pairs (F: 5′- GGGAAACGACAATCTACAGCTGAGAAGGTCAAGGC −3′ and R: 5′- TCAATATCATCTTCTTTTCCAGCGAAAACACCAGC −3′; F2: 5′- TACAAAGCCTGCGAACCAGCGATCATCGACCTTGA −3′ and R2: 5′- GTGAATTCGAGCTCGAAGATGGAGGTGGATGGGGA −3′, respectively).

The basic structure of pAg and *nptII* cassette (P*trpC* [GenBank accession no. X02390], *nptII*, T*cgrA* [AFUA_8G02750] were amplified from pAg 1−3’-UTR of ARB_02021 by PCR with specific primer pairs (F: 5′-CGAGCTCGAATTCACTGGCC−3′ and R: 5′- AGATTGTCGTTTCCCGCCTT−3′; F2: 5′-AGAAGATGATATTGAAGGAGCA−3′ and R2: 5′-TTCGCAGGCTTTGTACTTT−3′, respectively).

These four amplified fragments were joined with the In-Fusion system (TaKaRa Bio).

A Tr*cdc42*-targeting vector, pAg1P*_ctr4_*Tr*cdc42*, was constructed as follows: Approximately

1.7 and 2.0 kb of the upstream and ORF + downstream fragments of Tr*cdc42* (TERG_04946) were amplified from *T. rubrum* total DNA by PCR with specific primer pairs. (F: 5′- CGCACTAGTGGGATTTGGAGTCAAGGCGA−3′ and R: 5′- CGCGGGCCCCTCTCTCCTGCTGGTCTCCT−3′; F2: 5′- TACAAAGCCTGCGAACATGGTTGTTGCTACTATCA −3′ and R2: 5′- GTGAATTCGAGCTCGGAACGGGACAGGTTCGACTT −3′, respectively).

The upstream fragment and pAg1-*hph2*^72^ were cleaved with SpeI and ApaI enzymes and ligated. The ORF + downstream fragment containing vector was amplified by PCR with a specific primer pair(F: 5′-CGAGCTCGAATTCACTGGCC−3′ and R2: 5′-TTCGCAGGCTTTGTACTTT−3′, respectively). The ORF + downstream fragment and the linearized vector fragment containing upstream fragment were joined with the In- Fusion system (TaKaRa Bio). The PCRs were performed using Tks Gflex DNA polymerase (TaKaRa Bio).

### Recombinant protein preparation

TrRac, TrCdc42-His, and TrCdc24 DH-PH proteins were expressed using *E. coli* BL21 carrying pGEX6p-1 derivative plasmids. *E. coli* BL21 carrying the plasmid was precultured in 5 ml of LB with 50 μg/ml ampicillin at 37L. The preculture was then inoculated into 2 L of LB with 50 μg/ml ampicillin and cultured at 37°C for 3 h. Expression was induced with 0.1 mM IPTG, and the culture was maintained overnight at 20°C. During GST-TrCdc42-His induction, 4% ethanol was added. The purification of GST fusion proteins followed the instructions of GSTrap HP (GE Healthcare Japan, Japan). The purification buffer (Buffer A) consisted of 20 mM Tris/HCl (pH 7.5), 150 mM NaCl, 2.5 mM MgCl2, and 0.5 mM DTT. Purified fusion proteins were treated with PreScission protease at 4°C, and the flow-through of GSTrap was collected. After PreScission protease treatment, TrRac protein was desalted using Amicon UltraCel 10K (Merck, Germany), and the flow-through of Q sepharose HP (GE Healthcare Japan, Japan) was recovered. Following pre-scission protease treatment, Cdc42-His protein was further purified using Ni-NTA resin (FUJIFILM Wako Chemicals, Japan) and desalted using Amicon UltraCel10K (Merck, Germany). The purity of the TrRac, TrCdc42-His, and TrCdc24 DH-PH proteins was analyzed using fluorescence staining and stain-free imaging technology from Bio-Rad Laboratories (USA).

### Guanine nucleotide exchange assay

One μM of TrRac or TrCdc42-His was mixed with 0.1 μM Bodipy-FL GTP (Thermo Fisher, USA) in Buffer A and allowed to equilibrate for 3 min at room temperature. Ammonium sulfate, EDTA, or recombinant TrCdc24 DH-PH proteins were incubated with the compounds or vehicles alone (final DMSO concentration: 1%) at the indicated concentrations in the reaction buffer for 20 min at room temperature. The reaction was initiated by combining TrRac or TrCdc42/Bodipy-FL GTP (100 μl) with Cdc24DH-PH (50 μl) at room temperature. The change in Bodipy-FL-GTP fluorescence (excitation: 485 nm, emission: 535 nm) was monitored for 60 min using a Berthold tristar2 LB942.

### Transformation of *T. rubrum*

*T. rubrum* was transformed using the *A. tumefaciens* mediated transformation (ATMT) method as previously described with minor modifications.^10,69,73^ After cocultivation, nylon membranes were transferred onto Sabouraud dextrose agar (SDA) medium containing 200 μg/ml G418 (Nacalai Tesque, Japan), chloramphenicol, and 200 μg/ml cefotaxime sodium. They were then overlaid with 10 ml of SDA supplemented with 400 μg/ml of G418, chloramphenicol, and 200 μg/ml cefotaxime sodium and incubated for 7 to 14 days. Transformants were subsequently transferred onto new SDA plates with appropriate antibiotics. The desired transformants were screened by PCR and Southern blot analysis. Total DNA was extracted using the Quick-DNA Fungal/Bacterial Miniprep Kit (Zymo Research, USA). Beads beating of fungal cells was performed using μT-01 (TAITEC, Japan) with 5 mm stainless steel beads. Aliquots of approximately 0.5 - 1 μg of the total DNA were digested with an appropriate restriction enzyme, separated by electrophoresis on 0.8% (w/v) agarose gels, and transferred to Hybond-N+ membranes (GE Healthcare Ltd.). Southern hybridization was carried out using the ECL Direct Nucleic Acid Labeling and Detection System (GE Healthcare Ltd.) following the manufacturer’s instructions.

### Quantitative PCR

*T. rubrum* was precultured for one week in MOPS-buffered RPMI. The preculture was inoculated in MOPS-buffered RPMI with the desired concentration of CuSO_4_ or Bathocuproinedisulfonic acid (BCS). Harvested mycelium was homogenized using a bead shocker (TITEC, Japan). Total RNA was purified using the NucleoSpin RNA (MACHEREY-NAGEL, Germany). Reverse transcription and PCR reactions were conducted as previously described.^74,75^ For the quantification of Tr*cdc24* cDNA, the following primer pairs were used: F: 5′- ACAGAACCGGTACACTGCCC-3′ and R: 5′- AACGGAGGTAATGAGGGCCG -3′. For the quantification of Tr*cdc42* cDNA, the following primer pair was used; F: 5′- TGGAGATGAGCCATACACGC -3′ and R: 5′- TCTCAAAGGAAGCTGGCGAG -3′. The *rpb2* gene was used as an internal calibrator (primer: F: 5′- TGCAGGAGCTGGTGGAAGA −3′ and R: 5′-GCTGGGAGGTACTGTTTGATCAA−3′).^76^

### Cell morphological analysis

The cells were observed using a fluorescence microscope, specifically the BX53 Japan. Subsequently, the measurements of cell length and width were carried out utilizing Image J software, NIH. The polarity index was calculated as the length, which was measured at the longest diameter of a cell, divided by the width, measured at the midpoint of the length and perpendicular to the longest diameter, following the methodology outlined in reference.^78^

### Indirect fluorescence microscopy

The P*_ctr4_*Tr*cdc24* strain was inoculated with 1–5 × 10^6^ spores on sterile coverslips positioned in a 12-well plate. Subsequently, they were incubated with 500 μL of SD liquid medium overnight at 28°C. On the second day, the SD medium was refreshed with a fresh medium, and the spores were incubated once again overnight at 28°C. On the third day, the supernatant was carefully removed, and the cells were fixed with 4% paraformaldehyde (PFA, sourced from Nacalai Tesque, Japan) for 15 minutes at room temperature. Following this, the samples were washed three times with PBST (PBS + 0.05% Tween 20) and then incubated in 400 µL of 10 mg/ml lysing enzyme/10% bovine serum albumin (BSA)/PBS for 2 h at 28°C. Afterward, they were washed three times with PBST, permeabilized with 400 μL of pre-cooled methanol at −25°C for 10 min, and then washed again with PBST. The samples underwent incubation with blocking buffer (10% donkey serum/0.2% Triton X-100/0.02% sodium azide/PBS) for 30 min, followed by incubation with anti-beta-actin antibody (clone C-4, Merck; diluted 1/1000) in Can Get Signal A solution (Toyobo, Japan) at 4°C overnight. Following three washes with PBST, the samples were incubated with anti-mouse IgG antibody conjugated with Alexa488 (Abcam; diluted 1/1000) and DAPI solution (Dojindo, Japan; diluted 1/100000) in Can Get Signal A solution for 1 hour at room temperature. After three additional washes with PBST, the samples were rinsed with water, mounted on glass slides using Aqua-Poly/Mount (Polysciences, UK), and finally observed under a BZ-8100 all-in-one fluorescence microscope (Keyence, Japan) or an AX confocal microscope system (Nikon).

### Phylogenetic tree analysis

The evolutionary history was deduced using the neighbor-joining method.^77^The percentage of replicate trees in which the associated taxa clustered together in the bootstrap test (500 replicates) is depicted adjacent to the branches. Evolutionary distances were computed using the Poisson correction method and are presented as the number of amino acid substitutions per site. Positions containing gaps and missing data were excluded using the complete deletion option. The evolutionary analyses were carried out using MEGA X.^78^

### Invertebrate infection model of dermatophytosis

Eggs from silkworms were procured from Ehime-Sansh Co., Ltd. (Japan) and reared following a method described previously.^79,80^ Fifth instar silkworm larvae were nourished with an artificial diet (Silkmate 2S; Ehime-Sanshu Co., Ltd., Japan) overnight.^81,82^ The dermatophyte infection was conducted in accordance with a previously outlined procedure ^50^ Conidia (5 × 10^6^) of *T. rubrum* were injected into the silkworm hemolymph.

### *In vitro* nail infection assay

The conidial suspension (1 × 10^4^/10 μl/nail fragment) of *T. rubrum* was administered to autoclaved nail fragments obtained from an adult male volunteer (ethics approval, No. R3-6, Musashino University, Japan) and incubated at 28°C. Following 24 h period, either the vehicle (30% DMSO/5% Tween80) or 60 mM EHop-016 was applied to the nail fragment. A total of four applications were administered every 24 h. The nails were examined after 36 days of incubation.

### Statistical analysis

Statistical analyses were conducted using Microsoft Excel and R software. All experiments were performed more than three times. The determination of statistical significance was carried out using the Student’s *t*-test or one-way ANOVA-Tukey-Kramer. IC_50_ values were ascertained by plotting the fluorescent intensity (% of control) at 60 min using a specific formula.

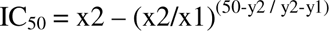

x1: a sample concentration at less than IC_50_,

x2: a sample concentration at higher than IC_50_

y1: fluorescence intensity (%) at x1,

y2: fluorescence intensity (%) at x2

