## Supplementary material for "Targeting dermatophyte Cdc42 and Rac GTPase signaling to hinder hyphal elongation and virulence": Figure S1

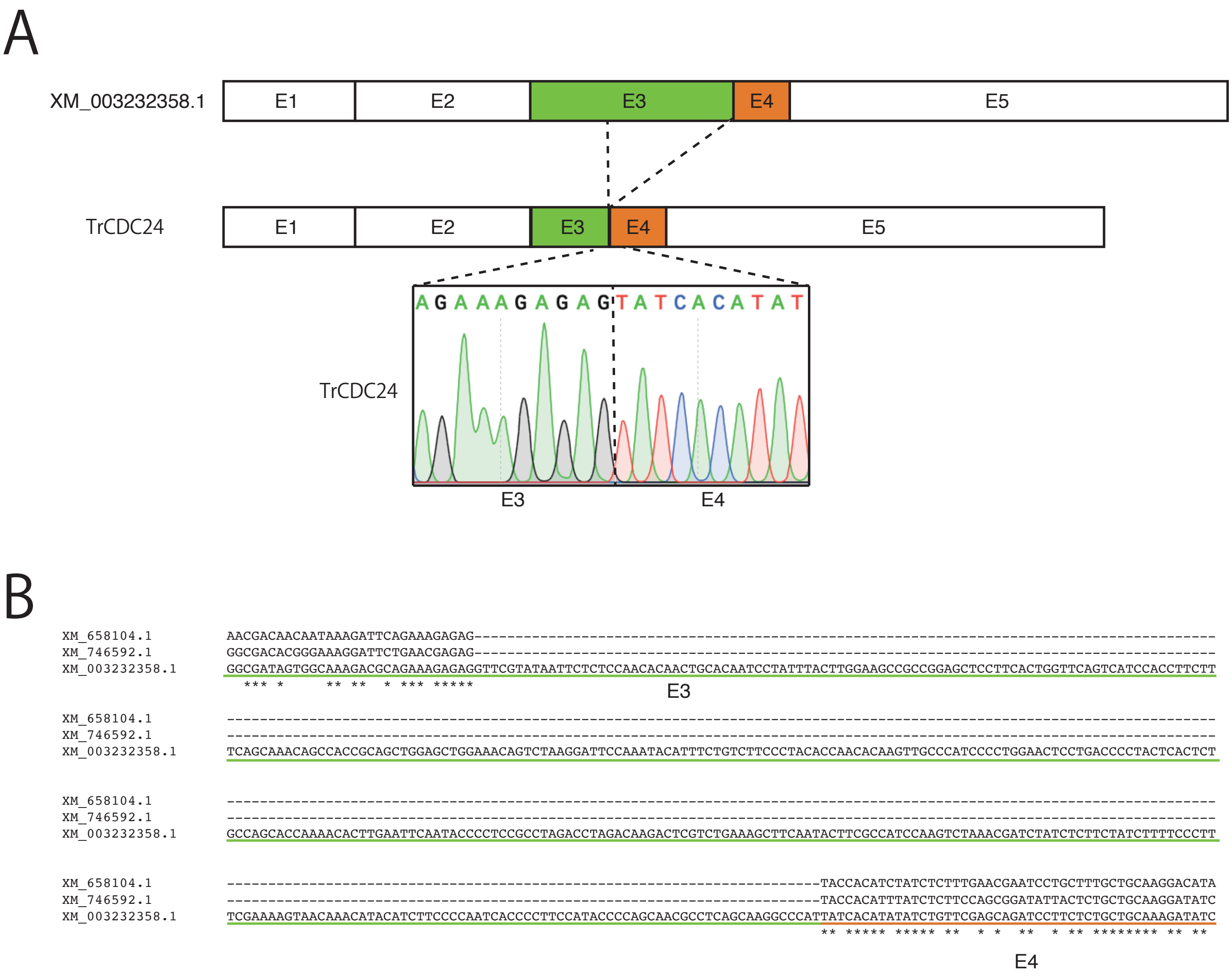


### Figure S1

**Figure S1 Part of XP_003232406.1 is a TrCdc42/TrRac GEF TrCdc24.**

A. Exons of XM_003232358.1 coding XP_003232406 and TrCdc24.

B. Partial sequences of *cdc24* gene in *A. nidulans* (XM_658104.1), *A. fumigatus* (XM_746592.1) and XM_003232358.1 exon 3 and exon 4 in *T. rubrum*.
